# Discrete protein dynamics enable long-range communication

**DOI:** 10.1101/2025.10.08.681220

**Authors:** Alan J. Katz, Levent Sari, Leigh J. Manley, Erdal Toprak, Milo M. Lin

## Abstract

Allostery, in which perturbations at an input protein site tune the activity at a distant output site, allows proteins to serve as molecular logic gates. Often, information is transmitted without altering the structure outside of the input and output sites. This focalized allostery requires correlated motion between protein backbone dihedral angles that are separated by distances many times longer than the scale of electrostatic interactions. What physical properties of folded proteins enable such long-distance information sharing despite thermal noise is unclear. To address this question, we introduce a Variable-Well Dihedral (VWD) model Hamiltonian which removes dependence on chemical details and forces, instead only tuning the degree of nonlinearity of purely local interactions within a densely-packed polymer. We show that tuning the physical parameters of the model gives rise to focalized allostery in so far that doing so increases the discreteness of the internal degrees of freedom, with real proteins occupying the highly discrete regime. These results parallel, at the molecular scale, the superiority of digital compared to analog signal processing for telecommuncations under noisy conditions.

## Introduction

Proteins can detect angstrom-scale chemical signals and robustly relay such perturbations across their nanometer-scale bodies. This propagation of information, termed allostery (*1*), is crucial to the molecular basis of information processing in living systems. For example, binding of a target molecule at the active site may occur only when a ligand binds to the allosteric site and induces a conformational change at the active site (Figure 1A).

**Figure 1:**
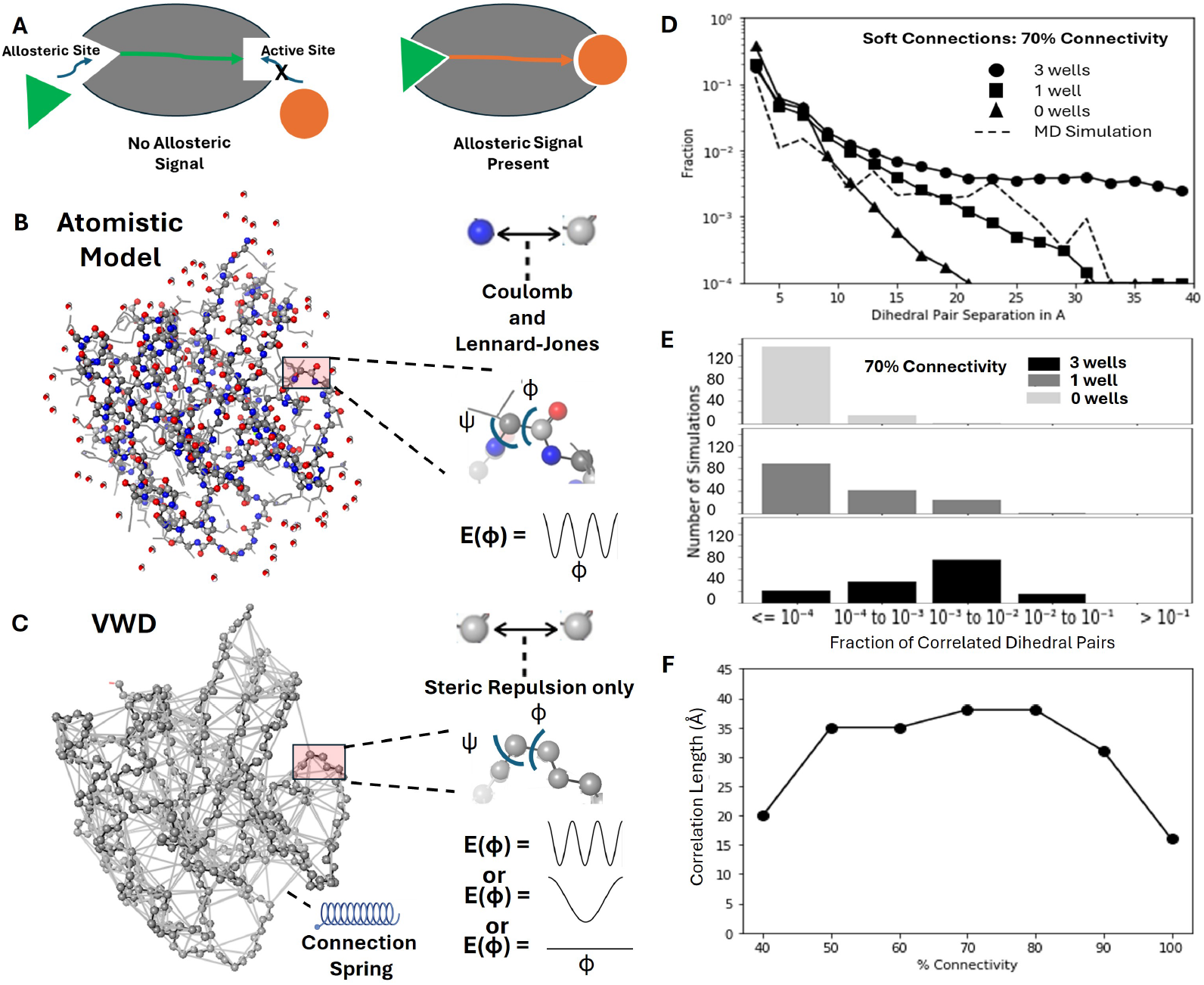
(A) Example of protein allostery: a ligand (orange circle) is unable to bind at the active site until a second ligand molecule (green triangle) binds at the allosteric site and triggers a conformational change at the active site. (B) A typical atomistic model used in molecular dynamics (MD) simulations shown for the HRAS protein (PDB ID 5P21). The atomistic model includes all atoms in the protein solvated by water and ion molecules, and non-bonded interactions between atom pairs arising from Coulomb and Lennard-Jones forces. The number of wells in the diheral angle potential energy function is 3 due to the nature of the covalent single bond. (C) In the Variable-Well Dihedral (VWD) model, only undifferentiated backbone atoms are defined, eliminating all side chains. Interatomic forces consist of a purely steric repulsion, and a network of harmonic springs (in light gray) serve to maintain the intramolecular network of contacts. As in the atomistic model, dihedral angles serve as the degrees of freedom (insets). In contrast to the atomistic model the VWD model allows for varying the number of dihedral potential wells (0, 1, or 3). (D) Simulation results for an ensemble of VWD simulations with variable set of randomly-chosen contacts initiated from the HRAS fold. The mean fraction of correlated dihedral pairs with significant mutual information calculated across the ensemble of VWD runs is shown as a function of the distance separating the pair. (E) Number of correlated dihedral pairs separated by at least 20 *Å* for the ensemble of runs used in D. (F) Correlation length in *Å* as a function of connectivity for ensembles of soft VWDs with 3 wells. Correlation lengths signify longest separation between dihedral pairs that share significant mutual information.

Allostery often involves only subtle changes in overall protein structure; the ligand-bound and ligand-free structures may be indistinguishable except for changes to structure or dynamics that are localized to the allosteric and active sites (*2–5*). Such focalized allostery is not explained by current models based on single- and multistate mechanisms (*6–13*) that result in global changes in protein structure. Atomistic simulations of protein dynamics can accurately replicate protein folding and dynamics, but their interactions are too detailed (Figure 1B) to allow systematic isolation of key physical constraints underlying allostery. Existing minimal models are typically 2D or 3D elastic network models (*10–13*) that can incorporate some aspects of protein physics by, for example, generating frustration via randomizing the rest lengths of the springs in the network (*11*). However, such models do not explicitly account for the nonlinear degrees of freedom and packing constraints characteristic of folded proteins, and which may be necessary for focalized allostery.

Focalized allostery manifests itself as mutual information (MI) between degrees of freedom that are separated by distances on the order of the protein size. Within proteins, these degrees of freedom are the dihedral angles of rotation about single covalent bond axes, which determine protein structure (see *ϕ* and *ψ* angles determining polymer geometry in Figure 1B). Molecular dynamics simulations of proteins at atomistic resolution show evidence of this long-range information transfer in the form of statistical correlations between discrete configurational states of dihedral angles in the active and allosteric sites (*14–16*). Allosteric networks based on these calculated correlations accurately predict allosteric sensitivity in a saturation mutagene-sis experiment (*17*). Yet, the physical interactions or constraints that enable such long-range information networks are unknown. To understand the necessary and sufficient conditions for creating focalized allostery, we introduce a Variable-Well Dihedral (VWD) model of a self-interacting folded polymer that enables explicit tuning of the degree of dihedral discreteness (or “digitalness”). Over a range of interaction densities, the model shows that dihedral discreteness is necessary and sufficient for focalized allostery. We run simulations of the VWD model with Langevin dynamics and find that long-range mutual information generally persists for different folding topologies and under physiologically relevant conditions of temperature and interaction density without parameter tuning, in contrast to existing elastic network models of protein allostery (*18*). When dihedral states are not discrete, long-range mutual information is suppressed by thermal noise. The link between the presence of focalized allostery in proteins and the discrete nature of dihedral states recalls the superior performance of digital over analog communication in noisy channels (*19*).

## Results

Our Variable-Well Dihedral (VWD) model provides a coarse-grained framework for modeling allostery that preserves constraints (*20, 21*) generic to proteins while removing chemical details that may act as confounders. This enables systematic interrogation of physical properties such as packing density and nonlinearity of local interactions that are necessary and sufficient for focalized allostery, while also allowing direct comparison with correlation lengths derived from atomistic simulations. The VWD model retains only the atoms that make up the backbone chain and eliminates side chains and backbone oxygens and hydrogens. The chemical identity of atoms along the backbone are removed such that all atoms are identical spheres (Figure 1C). Retaining only the backbone atoms allows us to define dihedral angles that parallel those found in the full protein. Whereas the complete model (Figure 1B) includes a full complement of long-range forces such as Lennard-Jones, the VWD model includes only a repulsive Lennard-Jones force to prevent atoms from passing through one another.

Both the atomistic model and the VWD model include a bare nonlinear dihedral rotation potential of the general form:

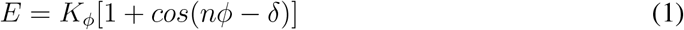

For the atomistic models (Figure 1B), the periodicity (n) is generally taken to be 3 for backbone *ϕ* and *ψ* dihedral angles, which corresponds to 0, 120, and 240 degrees, with no phase shift (*δ* = 0). In contrast, the VWD model allows for changes to the number of dihedral bare potential wells (Figure 1C) and randomly assigns values of *δ* to introduce frustration, leading to a degeneracy of low-energy protein configurations. K_*ϕ*_ energies are set to 2.5 kcal/mole (kT = 0.5922 kcal/mole at T = 298 K) and the *ω* and *ψ* dihedrals are fixed by assigning them energy barriers well above thermal energies.

Because the VWD model eliminates atomic identities, interactions, and side chain atoms, whose interactions are essential to determine the secondary and tertiary protein structure, we introduce a network of stabilizing connections between atoms familiar from elastic network models of proteins (*22*). To establish results that are not idiosyncratic to specific protein geometries or interaction patterns, we applied our model to an ensemble of protein-like structures. The ensemble is generated from the experimentally determined backbone atomic coordinates of a starting protein. A contact matrix is generated in which all distances between pairs of backbone C*α* atoms within a threshold distance of 7.3 *Å* are set to one (potential connected pairs), and all other entries are set to zero (unconnected pairs). Starting from this contact matrix, each member of the ensemble is generated by randomly choosing a given fraction of the connected pairs, with each chosen pair having the harmonic potential *V* = *K*(*r*–*r*_0_)^2^ with rest length *r*_0_ equal to the interatomic distance and spring constant *K* = 1*kcal/Å*^2^; we denote such connections as ‘soft’ because their energies are on scale of thermal fluctuations. This creates an ensemble of different alternate ‘protein folds’, each stabilized by a different randomly selected subset of spring connections from the contact matrix.

For each parameter set (different values of fractional contacts, spring constant, and dihedral potential), we generated on the order of 150 folds per ensemble and performed 24 microseconds of Langevin dynamics for each fold. We run dynamics simulations of the VWD model using the LAMMPS package (*23*), which allows considerable flexibility in setting potentials and force constants. Langevin dynamics is performed for all runs so that results include the impact of thermal noise while sampling conformational states according to equilibrium statistics in the canonical ensemble. In this study, we chose the starting protein to be H-Ras (PDB id 5p21 (*24*)), a 166 amino acid protein which plays a key signaling role in cell proliferation. We find considerable variation in the level and range of long-range mutual information within ensembles, hence, we report results in terms of statistics over each ensemble.

Figure 1D plots the mean fraction of dihedral pairs with significant mutual information (MI) versus the dihedral-pair separation distance for an ensemble in which 70% of the contact matrix are connected by springs. Significant MI is determined by a noise model based on atomistic MD simulations for a number of different proteins (*16*). Our noise model also accounts for spurious MI due to undersampling by subtracting from the count of dihedral pairs with significant MI instances following random translations of dihedral trajectories. The dashed curve denotes the results of MD simulations for the complete atomistic H-Ras model (*16*). The MD simulation curve shows a dropoff in MI around 25 *Å*, which likely reflects impact of H-RAS-specific side-chain interactions on dihedral motion. As seen in the figure, significant long-range MI for dihedral pairs persists in the VWD with 3-well dihedral potentials to length scales comparable to the linear dimension of the protein (*>*30 *Å*), while in the 0- and 1-well potential models, significant MI drops off rapidly at shorter length scales.

As a measure of the extent of long-range mutual information, we define the correlation length of mutual information to be the largest dihedral pair separation over which the mean fraction of dihedral pairs with significant MI lies above a threshold. This threshold is set by the point where we find a rapid drop in significant MI between dihedral pairs in the H-RAS MD dotted line curve in Figure 1D, which is approximately 25 *Å*. The H-RAS MD curve provides a biologically relevant measure of mutual information to define the correlation length in our plots. Figure 1E contrasts the difference in the number of correlated dihedral pairs with significant MI for at least 20 *Å* separations for the parameters sets shown in Figure 1D. For the various structures tested in the ensemble of simulation runs, having 3 dihedral wells, as opposed to 0 or 1 wells, significantly increases the likelihood of finding correlated dihedrals at longer length scales. Figure 1F shows the correlation lengths for a range of spring connectivities for the 3-well soft connection case. For connectivity that was too low (≤ 40%) or too high (100%), we observed significant drop-offs in correlation length. At 100% connectivity, the retention of all connections overconstrains many of the C*α* atoms, especially in parts of the protein structure corresponding to secondary and tertiary structures, and correlations between mobile dihedrals are limited to short length scales. At 40% connectivity, the degrees of freedom are insufficiently coupled to observe long-range correlations between dihedrals.

To better understand the role that the network of contact spring interactions plays in the absence or presence of long-range MI in the VWD, we performed a series of simulation runs with the contact spring constant K = 100 kcal/mole-*Å*^2^ (which we denote as the “hard” minimal model), so that collective dihedral rotations cannot change contact distances. Figure 2A shows the results for hard connections with 50, 60, and 70% connectivity. The correlation length for 50% connectivity and 3-well potentials is approximately 31 *Å*, which is on the order of the correlation lengths (*>* 31 *Å*) for soft 3-well models with 50-90% connectivity. However, the correlation lengths for the hard model with 50% connectivity and 0-well and 1-well potentials (23 and 26 *Å*, respectively) are significantly greater than what we observe for the corresponding 0-well and 1-well cases with 50% connectivity in the soft model. These results suggest that realization of long-range dihedral correlations in the VWD does not require soft, displaceable connections but rather additional constraints to the assumed dihedral states provided by multi-well potentials and/or hard connections. On the other hand, when the system is overconstrained as, for example, in the hard model with 70% connectivity and 3-well potentials, there are few dihedral transitions and mutual information drops off from MI found in the 50% cases for separations *>* 5 *Å*. Results for the hard minimal model suggest that long-range MI requires tuning of the number of connections to ensure that there are sufficient numbers of mobile and appropriately constrained dihedral states. In the case of the soft connections, much less tuning of the connectivity network is required to ensure a population of mobile dihedral states, although a connectivity below 100% remains a pre-requisite for long-range MI (Figure 1F). These results indicate that the conditions for focalized allostery are robust to the details of the VWD model.

**Figure 2:**
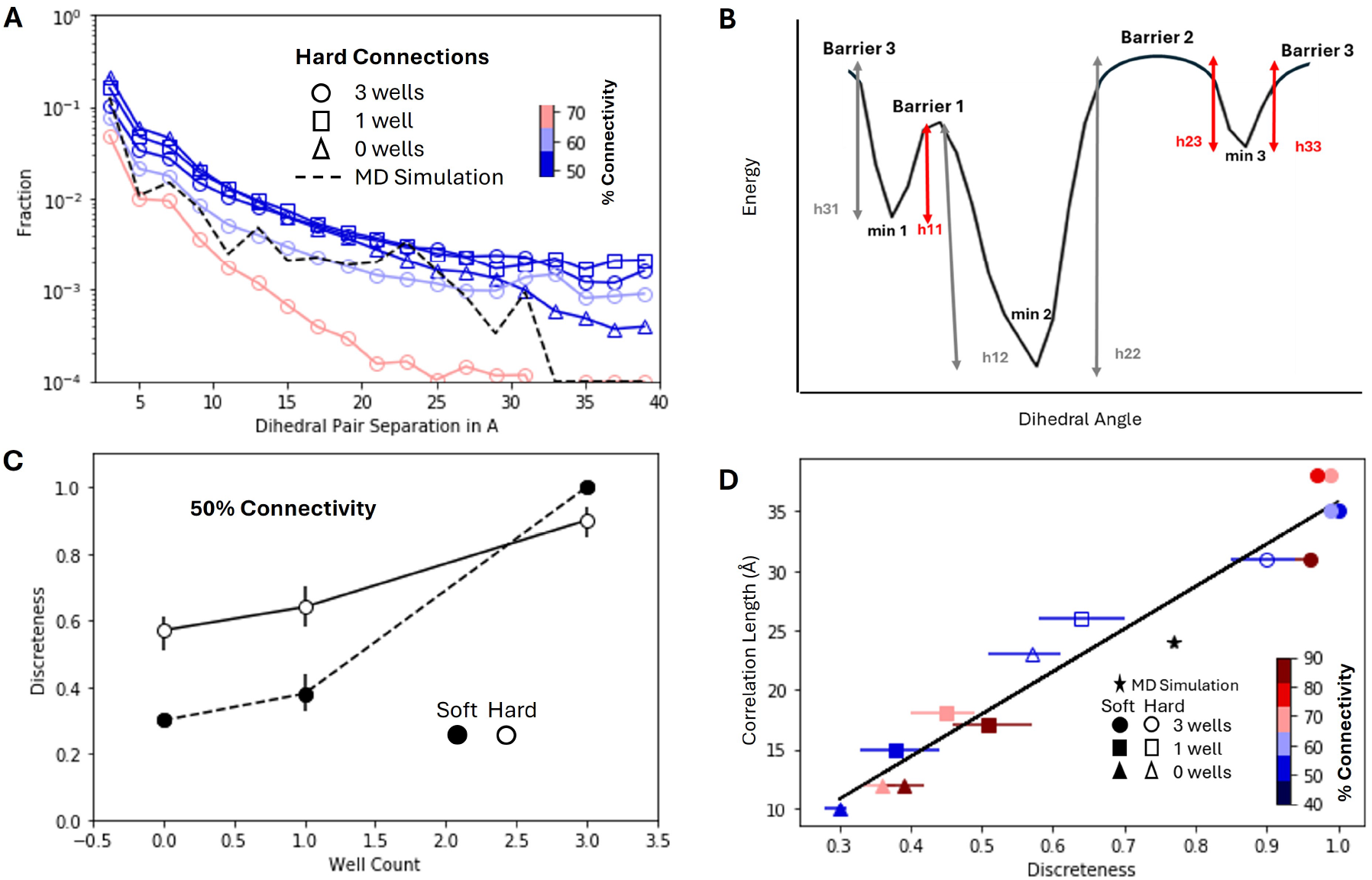
(A) Panel A shows simulation results for an ensemble of VWD runs with hard connections (spring constant K = 100 kcal/*Å*^2^) and 0,1, and 3 dihedral potential wells at 50% matrix connectivity and for 3 wells at 60% and 70% matrix connectivity. (B) Panel B illustrates calculation of discreteness for a multi-state dihedral. For each barrier *b*, we find its two barrier heights *hb*,m1[*b*] and *hb*,m2[*b*] between each of the two neighboring minima *m*1 and *m*2 on either side of the barrier; the minimum height associated with the barrier is defined to be *h*_*b,min*_ = min[*hb*,m1[*b*], *hb*,m2[*b*]]. The dihedral discreteness is then: ⟨tanh(*h*_b,min_)⟩_*b*_, where the minimum barrier height is passed through a tanh function to ensure that the discreteness falls between 0 and 1 and ⟨⟩_b_ denotes theaverage over all barriers of the dihedral. (C) ‘Well Count’ is the number of wells in the bare dihedral potential and ‘Discreteness’ is defined in Panel B and the text. (D) Correlation lengths versus median discreteness values for ensembles of the VWD for different matrix connectivity, numbers of dihedral potential wells, and soft and hard connections. Correlation length is the longest length scale over which significant long-range MI is observed between dihedral pairs. Horizontal error bars signify first quartile and third quartile values of discreteness for various ensembles of minimal model runs.

We introduce a measure of dihedral discreteness to capture how cleanly the dihedral angular states partition into distinguishable states for variations of the VWD model. Our measure captures how kinetically stable multistate dihedrals in a given protein structure are. To quantify discreteness, we first convert the dihedral angle probability function *P* into a free energy *G*: *P* = exp(-*G*/*kT*)/*Z*, where *Z* is the partition function. Only dihedrals with multiple minima in their free energy function have discreteness. Figure 2B illustrates how the discreteness is calculated for a given multi-state dihedral. For each barrier *b*(*i*) of multi-state dihedral *i*, we find its two barrier heights *h*_*b*(*i*),m1[*b*(*i*)]_ and *h*_*b*(*i*),m2[*b*(*i*)]_ between each of the two neighboring minima m1 and m2 on either side of the barrier; the minimum height associated with the barrier is defined to be *h*_*b*(*i*),*min*_ = min[*h*_*b*(*i*),m1[*b*(*i*)]_, *h*_*b*(*i*),m2[*b*(*i*)]_]. The discreteness of the entire protein is then:

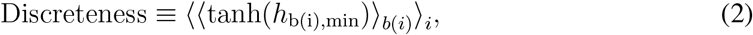

where the minimum barrier height is passed through a tanh function to ensure that the discreteness lies between 0 and 1, ⟨⟩_i_ denotes the average over all multi-state dihedrals *i* in the protein and ⟨⟩_b(i)_ denotes the average over all barriers of dihedral *i*.

Figure 2C shows how dihedral discreteness varies for the soft and hard VWD connections as a function of the number of wells in the dihedral energy function (Equation 1). There is a clear trend of increasing discreteness with the degree of imposed constraints that stem either from the number of bare potential wells or the hardness of connections. Moreover, the observed range for the hard VWD as a function of the well count is not as great as that for the soft VWD, since the added constraint provided by hard connections compensates for lower well count.

Figure 2D shows a relationship between discreteness and measured correlation lengths for soft and hard VWDs with various connectivity and number of bare potential wells. The x-axis is the median discreteness of the protein for a set of runs, and the error bars on the lower and upper x axes signify the first and third quartiles, respectively. The effects of all the different parameter choice combinations on the correlation length collapse onto a single variable: discreteness. Discreteness arises from the constraints imposed by multi-well dihedral potentials and hard connections, which confine dihedral motion to narrow ranges of angles and generally limit the number of states assumed by the dihedrals. High discreteness correlates with long-range MI for both hard- and soft-minimal models with 3-well dihedral potentials, and low discreteness corresponds to the absence of long-range MI in soft-minimal models with 0-well and 1-well potentials. Hard minimal models with 0- and 1-well potentials manifest intermediate levels of discreteness and long-range MI as does the MD simulation for H-RAS (indicated by the star in Figure 2D). As noted earlier, the lower values of MI at larger dihedral separations for the full-atomistic H-RAS model likely reflects additional constraints on dihedral motion from side-chain interactions. In general, the discreteness variable captures the impact of constraints on long-range MI regardless of whether the source constraints is the shape of the dihedral potential function or the stiffness of springs.

## Discussion

We introduce the VWD model for proteins that accounts for long-range mutual information observed in MD simulations and multiple proteins (*14,16*). The observed long-range mutual information reflects focalized allostery, which is shown to be linearly correlated with discreteness regardless of the details of the nonlinear model. This is consistent with the fact that dihedrals in proteins assume a small number of distinct angular states (*25*). The dihedral discreteness, which stems in the VWD from a combination of constraints owing to nonlinear dihedral potentials with multiple minima as well as interactomic distance constraints, is necessary and sufficient for focalized allostery under biologically relevant packing and interaction densities. The notion of discreteness suggests that information transfer in proteins shares the advantages digital signal processing systems have over analog systems, which are more susceptible to information loss under noisy conditions (*19*).

The scaffold of connections between C*α* atoms in the VWD defines regions of varying levels of connectivity and, therefore, differing degrees of rigidity. Generally, regions with higher connectivity and higher rigidity correspond to various features of the protein structure, such as *β* sheets, *α* helixes, and hydrophobic cores (*26*). Others (*27, 28*) have explored protein dynamics from the perspective of rigidity analysis. In such studies, rigid regions of the material move as single-bodies facilitated by hinge-like motion of atoms along the interface between the rigid bodies and the more flexible regions of the material. Motion of these rigid regions requires coordination of the hinges along the rigid-flexible interfaces, which, in our VWD, occurs through coordinated dihedral rotations.

Parallels exist between rigidity analysis and the coordination of dihedral motions observed in the hard VWD, where long-range mutual information is only observed over a limited range of matrix connectivity. For example, we can consider the hard VWD in the context of Maxwell’s criterion to stability in mechanical lattice structures (*29*), where the critical average coordination number Z_*c*_, which separates floppy from stable rigid frames in *d* spatial dimensions containing *N* lattice nodes is: *Z*_*c*_ = 2*d* − [*d*(*d* + 1)*/N*]. Using this criterion, we find that the hard VWD discussed in Figure 2A should first show floppy modes below a lattice connectivity of 60%. This is consistent with the appearance (absence) of long-range mutual information in the hard VWD at 50% (60% and 70%) connectivity.

Our results suggest that long-range mutual information is observed in the soft VWD over a significant range of matrix connectivity, as shown in Figure 1F. This means that the more physically relevant soft VWD is robust and does not have to be finely tuned to realize long-range mutual information and focalized allostery. Evolution may have found paths to optimize these allosteric features in real proteins, as their foundational usefulness to protein and organism function has developed over time. Evolutionary trial and error often leverages intrinsic physical properties to engineer novel capabilities: in this case, the ability of proteins to encode information in a discrete format enables long-range information transmission in noisy cellular environments.

## Acknowledgements

The authors would like to thank Madhave Mani and Efi Efrati for helpful feedback on this work. This work was supported by NIH grant R01GM125748 and CZI Theory Institute Without Walls (M.M.L.)

## Notes

### Competing Interest Statement

The authors have declared no competing interest.

